# Differential expression of an alternative splice variant of IL-12Rβ1 impacts early dissemination in the mouse and associates with disease outcome in both mouse and humans exposed to tuberculosis

**DOI:** 10.1101/271627

**Authors:** Mrinal K. Das, Aurelie A. Ray, Yi Cai, Akul Singhania, Christine M. Graham, Mingfeng Liao, Jeffrey J. Fountain, John E. Pearl, Manish Pareek, Pranab Haldar, Anne O’Garra, Xinchun Chen, Andrea M. Cooper

## Abstract

Experimental mouse models of TB suggest that early events in the lung impact immunity. Early events in the human lung in response to TB are difficult to probe and their impact on disease outcome is unknown. We have shown in mouse that a secreted alternatively-spliced variant of IL-12Rβ1, lacking the transmembrane domain and termed ΔTM-IL-12Rβ1, promotes dendritic cell migration to the draining lymph node, augments T cell activation and limits dissemination of *M. tuberculosis* (Mtb). We show here that CBA/J and C3H/HeJ mice (both highly susceptible to Mtb) express higher levels of ΔTM-IL-12Rβ1 than resistant C57BL6 mice and limit early dissemination of Mtb from the lungs. Both CD11c+ cells and T cells express ΔTM-IL-12Rβ1 in humans, and mice unable to make ΔTM-IL-12Rβ1 in either CD4 or CD11c expressing cells permit early dissemination from the lung. Analysis of publically available blood transcriptomes indicates that pulmonary TB is associated with high ΔTM-IL-12Rβ1 expression and that of all IL-12 related signals, the ΔTM-IL-12Rβ1 signal best predicts active disease. ΔTM-IL-12Rβ1 expression reflects the heterogeneity of latent TB infection and has the capacity to discriminate between latent and active disease. In a new Chinese TB patient cohort, ΔTM-IL-12Rβ1 effectively differentiates TB from latent TB, healthy controls and pneumonia patients. Finally, ΔTM-IL-12Rβ1 expression drops in drug-treated individuals in the UK and China where infection pressure is low. We propose that ΔTM-IL-12Rβ1 regulates early dissemination from the lung and that it has diagnostic potential and provides mechanistic insights into human TB.

## Introduction

Tuberculosis (TB) remains a critical worldwide health issue despite significant public health and research efforts^1^. Key elements in our failure to limit this disease are the large number of people infected with the bacterium, *Mycobacterium tuberculosis* (Mtb) which causes disease, the lack of an effective vaccine against pulmonary TB and the need for prolonged and multiple drug treatments^2^. A critical gap in knowledge is what separates those that develop disease from the majority of infected individuals who control the infection without consequences^3^. Mendelian susceptibility to mycobacterial disease (MSMD) provides some insight, as those lacking the ability to generate or respond to interferon (IFN)-γ have a significantly increased risk of mycobacterial disease including TB^4^. A key component identified by MSMD and one known to exacerbate severity of TB is genetic dysfunction of the Interleukin (IL)-12 receptor component IL-12Rβ1 ^4, 5^, which acts with IL-12Rβ2 or IL-23R to mediate signaling by IL-12p70 or IL-23 respectively^6^. Understanding how the IL-12 receptor complex contributes to protection and disease is therefore critical. While the major function of this receptor complex is to promote IFN-γ production and consequent macrophage activation^4, 6^, we have identified a novel role for IL-12p40 and IL-12Rβ1 in initiation of early events within the lung following low dose aerosol exposure to Mtb ^7, 8^.

Experimental mouse models of aerosol exposure to Mtb indicate that the rapidity of bacterial dissemination and early activation of T cells in the draining lymph node and spleen contribute to effective control of lung infection^9, 10, 11^. Specifically, slow dissemination of bacteria leads to delayed T cell activation and delayed recruitment of these T cells^12^. This delay results in T cells arriving at the infected foci after the inflammatory response to the bacteria has begun and where the inflamed environment limits the efficacy of the T cell response in situ ^10, 13^. We have shown that the cytokine IL-12p40 is expressed early following Mtb exposure ^8^ and that if dendritic cells do not express IL-12p40 then they do not become motile when exposed to Mtb, do not migrate from the lung to the draining lymph node and do not drive naive T cell activation ^7^. Dendritic cells in the lungs of Mtb-exposed mice produce not only IL-12p40 but also an alternative splice variant of IL-12Rβ1 which lacks the transmembrane domain and which we have called ΔTM-IL-12Rβ1 ^8^. This splice variant was identified when the receptor was first described^14^ however its function was not appreciated until our studies in Mtb^8, 15^. The ΔTM-IL-12Rβ1 locates to the endoplasmic reticulum^16^ and can be secreted by transfected cells and mitogen-activated T cells blasts^15^. Human cells produce a similar splice variant of the IL12RB1 gene in response to Mtb exposure and this is referred to as isoform 2 ^8, 17^ and has been shown to augment human T cell responsiveness to IL-12p70^18^. Our data supports the hypothesis that the alternative splice variant of IL-12Rβ1 plays an early role in initiation of T cell activation in response to aerosol infection with Mtb and that the production of this splice variant in human TB merits investigation.

We have used meta-analysis of public data sets with statistical analysis to combine results from independent but related studies to identify factors potentially contributing to human disease^19^. Meta-analysis of gene expression data in particular can provide support for further investigation of any one particular pathway by targeted differential expression analysis20, analysis of relative expression of gene in various tissues and cells^21^ and pathway analysis ^22^. Multiple transcriptomic profiles of TB patients and controls have been deposited for analysis^23^. These data sets are available from areas of the world where the chance of infection (or infection pressure) varies due to the incident rate of disease; thus South Africa has a high incidence of TB at 781/100,000 and thus has high infection pressure, China has moderate incidence at 65/100,000 (modest infection pressure) and the UK has an overall low incidence (10/100,000) with some higher areas in London and the Midlands but at best only a low infection pressure^1^. This variety in the source of transcriptomic data sets allows for global insight into how specific gene expression profiles are associated with disease and also allows infection pressure to be considered as a factor in this association. Importantly for our study, while the commercially available arrays have probes sets specific for IL12RB1 these sets contain probes uniquely specific for either the full length or the alternatively spliced isoform. Combining the signal for the full and the alternatively spliced isoform into one signal for IL12RB1 obfuscates which isoform is expressed and therefore manual curation is required in order to differentiate these signals.

To address the potential role of the alternative splice variant of IL-12Rβ1 in TB we have combined mouse mechanistic studies with analysis of both published and novel transcriptomic profiles. We have found that high expression of ΔTM-IL-12Rβ1 in the lungs of mice susceptible to Mtb results in very limited early dissemination of bacteria from the lung to the periphery. Furthermore, mice lacking the ability to make the ΔTM-IL-12Rβ1 in either CD11c^+^ or CD4^+^ expressing cells are compromised in their ability to limit dissemination to the periphery. Taken together these data implicate early expression of the ΔTM-IL-12Rβ1 as a factor in control of Mtb dissemination from the lung to the periphery. To determine if the human ΔTM-IL-12Rβ1 or isoform 2 is associated with infection and disease in areas of low and high infection pressure we analyzed expression of the ΔTM-IL-12Rβ1 specific probes in transcription data sets from around the world. We found that expression of ΔTM-IL-12Rβ1 in the blood is associated with active pulmonary TB and that it reflects the heterogeneity of the latently infected population.

## Materials and methods

### Subjects and clinical sample collection

Protocols were approved by the Institutional Review Board of Shenzhen Third People’s Hospital, Shenzhen, China and informed consent was obtained from all participants. Whole blood samples were collected from healthy controls (HC, n=20), individuals with latent tuberculosis infection (LTBI, n=20), patients with active tuberculosis (TB, n=20) and patients with pneumonia (n=20) at Shenzhen Third People’s Hospital.

Patients with TB were diagnosed based on clinical symptoms, chest radiography, positive sputum Mtb culture and positive response to anti-TB chemotherapy. Asymptomatic individuals with non-clinical disease were recruited as controls. Mtb specific IGRAs were used to differentiate individuals with LTBI from HCs without infection ^24^. The diagnosis of pneumonia was based on 1) lavage fluid or sputum cultures were Mtb-negative during clinical follow-up, 2) new infiltration and clinical signs on chest radiograph were evident and completely resolved after treatment with the appropriate antibiotics, and 3) viral pathogens were not detected. The demographic characteristics of study populations of this study are in Table 1.

**Table 1:** The demographic characteristics of study population Shenzhen Third Peoples Hospital in Shenzhen, China.

| Study Cohort | n | Median age in years (range) | Gender (M/F) | AFB-positive | ELISPOT Positive |
| --- | --- | --- | --- | --- | --- |
| TB | 20 | 38.2 (25.0-58.0) | 13/7 | 20 | 20 |
| LTBI | 20 | 31.2 (23.0-56.0) | 12/8 | ND<sup>1</sup> | 20 |
| HC | 20 | 28.6 (21.0-48.0) | 10/10 | ND | 0 |
| Pneumonia | 20 | 39.2 (29.0-59.0) | 12/8 | ND | ND |

<sup>1</sup>ND = not done

### Mice

C57BL/6J, C3H/HeJ and CBA/J mice were purchased from JAX mice and housed at Trudeau Institute, Inc. B6.Cg-Tg(CD4-Cre) and B6.Cg-Tg(Itgax-cre) mice were purchased from Jax and crossed with our transgenic mice, which upon FLP and CRE recombination excise introns 13 and 14 of the *il12rb1* gene^15^. Confirmation of excision was undertaken with PCR in each case.

### Infection

Mtb strain H37Rv was grown in Proskauer Beck medium containing 0.05% Tween 80 to mid-log phase and frozen at –70°C. Mice were infected with a low dose of bacteria (~75 CFU) using a Glas-Col airborne infection system as previously described^25^. At day 1 and day 15 post-challenge, mice were killed by CO_2_ asphyxiation, the organs were aseptically excised and individually homogenized in saline. Organ homogenates were subsequently plated on nutrient 7H^11^ agar (BD Biosciences) for 3 weeks at 37°C, at which point CFU were counted.

### Quantitative PCR

RNA was generated from mouse tissue or human cells using the RNAeasy kit and was reverse transcribed with an RT Kit from Life technologies – now ThermoFisher Scientific). RT-PCR was performed to generate Ct values which were normalized to the housekeeping gene signal and then applied to a standard curve to determine copy number for each isoform of human and mouse IL-12Rβ1 transcript generated from a specific template^26^.

### Human monocyte derived dendritic cells

Discarded filters from blood donor clinic were harvested for peripheral blood cells and the CD14+ cells were purified using magnetic beads (Miltenyi). CD14+ cells were seeded at 2×10^6^/ml into wells in RPMI containing 10% FCS and stimulated with 4000IU/ml GM-CSF and 1000IU/ml IL-4 (both cytokines from Peprotech). Media was refreshed at 3 days and the cells were used for experiments at day 5.

### Microarray data from publically available databases

To evaluate gene expression of ΔTM-IL-12Rβ1 in TB, we used the transcriptome-microarray datasets available in public repositories. Briefly, public databases of Gene Expression Omnibus (http://www.ncbi.nlm.nih.gov/geo) and ArrayExpress (https://www.ebi.ac.uk/arrayexpress) were searched with the keywords ‘Tuberculosis, Patients’. A total of 59 data series (58 in GEO and 19 in ArrayExpress – with 16 overlapping) were retrieved from the search. We used the following exclusion criteria:-samples other than blood or blood derived cells, array platform lacking specific probe for ΔTM-IL-12Rβ1, series lacking non-comorbid TB conditions, lacking adult TB subjects, with less than 3 replicated samples for one condition and repeat arrays, we identified 16 arrays for further analysis. Full texts of the associated articles were evaluated for details on patient selection, demographics, sampling and RNA analysis. Six out of 16 were further excluded as the median centered across samples was not zero ^27^ and final analysis was carried out on 10 datasets (Table 2). *Processing of microarray dataset*. Out of 10 data sets, 7 were generated on Illumina and 3 were on the Affymetrix platform (Table 2). All 3 Affymetrix data series were generated on the same platform. The raw.cel files were converted to expression values using GC-RMA package read by ‘affy’ package on Bioconductor^21^. Associated packages and libraries used were “hgu133plus2.db”, “hgu133plus2cdf”. R-studio version-1.0.143.0 using Bioconductor libraries and R statistical packages were used for data analysis. Processed data matrices were exported into tab delimited txt files.

**Table 2.**
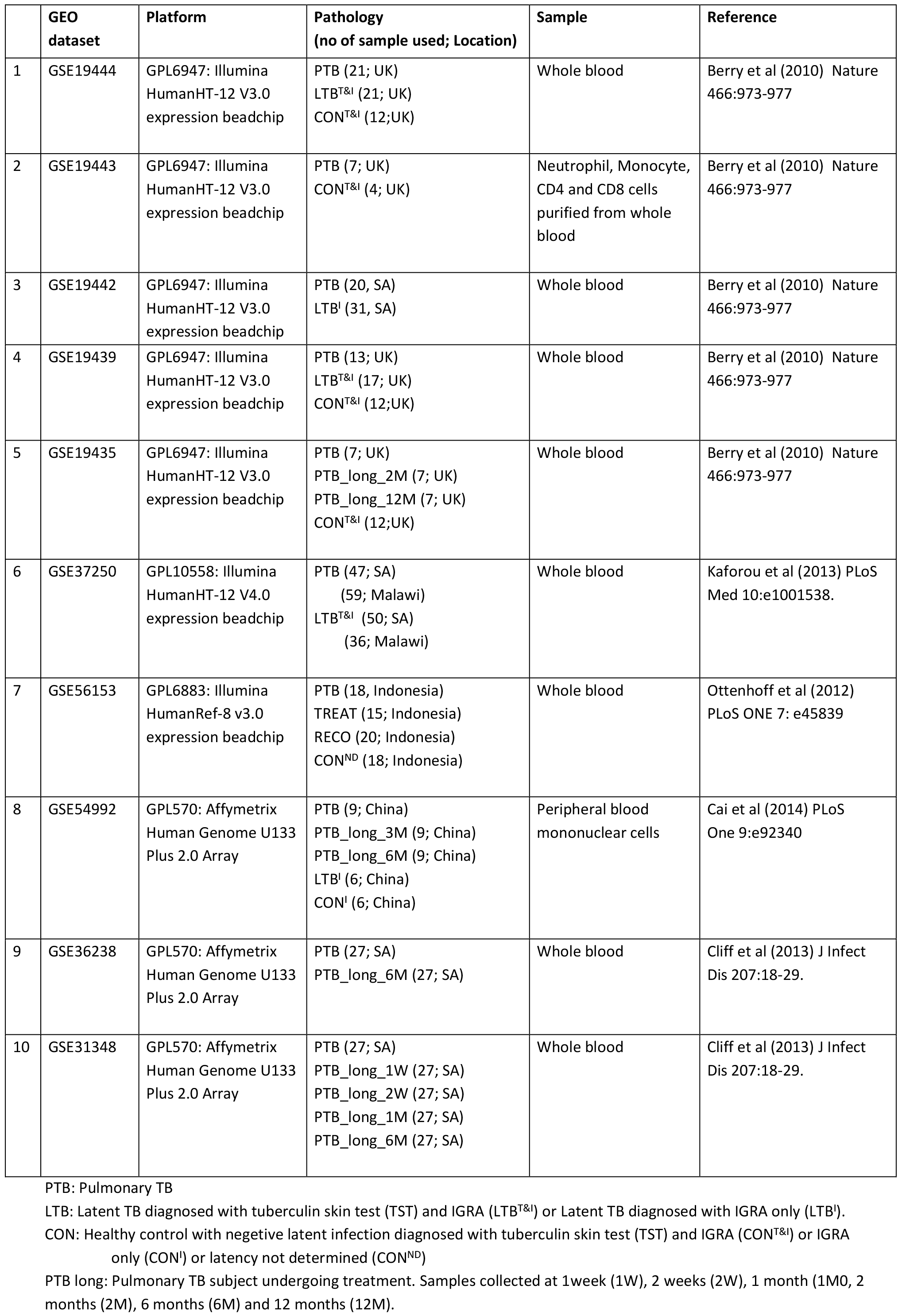
Details of the public array data assessed for expression of the ΔTM-IL-12Rβ1 sequence in blood samples from individuals with pulmonary TB, latent TB or other diseases.

|  | GEO dataset | Platform | Pathology<br>(no of sample used; Location) | Sample | Reference |
| --- | --- | --- | --- | --- | --- |
| 1 | GSE19444 | GPL6947: Illumina HumanHT-12 V3.0 expression beadchip | PTB (21; UK)<br>LTB<sup>T&I</sup> (21; UK)<br>CON<sup>T&I</sup> (12;UK) | Whole blood | Berry et al (2010) Nature 466:973-977 |
| 2 | GSE19443 | GPL6947: Illumina HumanHT-12 V3.0 expression beadchip | PTB (7; UK)<br>CON<sup>T&I</sup> (4; UK) | Neutrophil, Monocyte, CD4 and CD8 cells purified from whole blood | Berry et al (2010) Nature 466:973-977 |
| 3 | GSE19442 | GPL6947: Illumina HumanHT-12 V3.0 expression beadchip | PTB (20, SA)<br>LTB<sup>I</sup> (31, SA) | Whole blood | Berry et al (2010) Nature 466:973-977 |
| 4 | GSE19439 | GPL6947: Illumina HumanHT-12 V3.0 expression beadchip | PTB (13; UK)<br>LTB<sup>T&I</sup> (17; UK)<br>CON<sup>T&I</sup> (12;UK) | Whole blood | Berry et al (2010) Nature 466:973-977 |
| 5 | GSE19435 | GPL6947: Illumina HumanHT-12 V3.0 expression beadchip | PTB (7; UK)<br>PTB_long_2M (7; UK)<br>PTB_long_12M (7; UK)<br>CON<sup>T&I</sup> (12;UK) | Whole blood | Berry et al (2010) Nature 466:973-977 |
| 6 | GSE37250 | GPL10558: Illumina HumanHT-12 V4.0 expression beadchip | PTB (47; SA)<br>(59; Malawi)<br>LTB<sup>T&I</sup> (50; SA)<br>(36; Malawi) | Whole blood | Kaforou et al (2013) PLoS Med 10:e1001538. |
| 7 | GSE56153 | GPL6883: Illumina HumanRef-8 v3.0 expression beadchip | PTB (18, Indonesia)<br>TREAT (15; Indonesia)<br>RECO (20; Indonesia)<br>CON<sup>ND</sup> (18; Indonesia) | Whole blood | Ottenhoff et al (2012) PLoS ONE 7: e45839 |
| 8 | GSE54992 | GPL570: Affymetrix Human Genome U133 Plus 2.0 Array | PTB (9; China)<br>PTB_long_3M (9; China)<br>PTB_long_6M (9; China)<br>LTB<sup>I</sup> (6; China)<br>CON<sup>I</sup> (6; China) | Peripheral blood mononuclear cells | Cai et al (2014) PLoS One 9:e92340 |
| 9 | GSE36238 | GPL570: Affymetrix Human Genome U133 Plus 2.0 Array | PTB (27; SA)<br>PTB_long_6M (27; SA) | Whole blood | Cliff et al (2013) J Infect Dis 207:18-29. |
| 10 | GSE31348 | GPL570: Affymetrix Human Genome U133 Plus 2.0 Array | PTB (27; SA)<br>PTB_long_1W (27; SA)<br>PTB_long_2W (27; SA)<br>PTB_long_1M (27; SA)<br>PTB_long_6M (27; SA) | Whole blood | Cliff et al (2013) J Infect Dis 207:18-29. |

PTB: Pulmonary TB
LTB: Latent TB diagnosed with tuberculin skin test (TST) and IGRA (LTB<sup>T&I</sup>) or Latent TB diagnosed with IGRA only (LTB<sup>I</sup>).
CON: Healthy control with negative latent infection diagnosed with tuberculin skin test (TST) and IGRA (CON<sup>T&I</sup>) or IGRA only (CON<sup>I</sup>) or latency not determined (CON<sup>ND</sup>)
PTB long: Pulmonary TB subject undergoing treatment. Samples collected at 1week (1W), 2 weeks (2W), 1 month (1M), 2 months (2M), 6 months (6M) and 12 months (12M).

For Illumina data series matrix, those including non-log transformed and normalized data were analyzed using ‘GEOquery’ and ‘lumi’ package^28, 29^. For datasets having transformed and normalized values in series matrix, raw (non-normalized data) data files were downloaded from the GEO_supplementary files followed by processing using ‘lumi’ package. Briefly, data sets were processed, background corrected, log-2 transformed and then normalized using ‘quantile’ method. The processed data file was saved as tab delimited .txt file.

### RNA-seq data analysis

Raw paired-end RNA-seq data from Singhania et al. ^30^ for the Berry London cohort^31^ was subjected to quality control using FastQC (Babraham Bioinformatics) and MultiQC^32^. Trimmomatic v0.36 was used to remove adapters and filter raw reads below the 36 bases long and leading and trailing bases below quality 25^33^. Paired filtered reads from Trimmomatic were aligned to the *Homo sapiens* genome Ensembl GRCh38 (release 86) using HISAT2 v2.0.4 with default settings and RF rna-strandedness^34^. Gene transcripts representing multiple splice variants for each gene locus were assembled using StringTie v1.3.3 with default settings, using the-eB and stranded library fr-secondstrand parameters, and the reference genome mentioned above^35^. The assembled transcriptomes were annotated and analyzed using the *bioconductor* package Ballgown^36^ in R to obtain fragments per kilobase of transcript per million reads sequenced (FPKM) values^36^.

### QuantiGene RNA analysis

Whole blood was collected directly into PAXgene Blood RNA tubes (BD Biosciences) from individuals classified as HC, LTBI, TB and pneumonia in Shenzhen’s Third People’s Hospital, China (Table 1). Total RNA was extracted using the PAXgene Blood RNA Kit (Qiagen) following the manufacturer’s instructions. The purified RNA samples were processed and run on a custom Panomics QuantiGene 2.0 Multiplex array (Affymetrix) plate by the Shanghai Biotechnology Corporation (China, Shanghai). This plate was designed by the manufacturer with probes to simultaneously detect 31 target genes including IL12RB1. Gene expression was normalized to Housekeeping genes transcripts following the manufacturer’s protocol (http://www.panomics.com).

### Statistical analysis

Differences between the means of experimental groups were analyzed using the two tailed Student’s *t*-test or ANOVA as appropriate. Paired *t* tests were used for the longitudinal data. Differences were considered significant where *P*≤ 0.05. Inherently logarithmic data was transformed for statistical analysis. Where bacterial numbers were below the limit of detection the samples were given the value of the limit of detection to allow analysis. Pearson correlation was used to assess interaction between probes within a dataset. Briefly lumi/GC-RMA processed intensity data were used to prepare a functional data matrix with selected variables, this data matrix was assessed in the ‘Statistical Module” of the Metaboanalyst 3.0 online tool to calculate the Pearson Correlation Coefficient^37^. Similarly, a modified data matrix with downselected variables was used for Receiver Operating Characteristic (ROC) curve analysis. The analysis was carried out in ‘Biomarker Analysis’ module of MetaboAnalyst 3.0 online tool^37^. Classical univariate ROC curve analysis was used for calculation of the area under curve (AUC).

## Results

### Expression of ΔTM-IL-12Rβ1 in the lung associates with reduced dissemination of Mtb from the lung to the periphery

We knew from our previous studies that the ΔTM-IL-12Rβ1 message was induced early in the lungs of C57BL/6J (B6) mice exposed to low dose aerosol infection^8^. We also knew that the global inability to make the ΔTM-IL-12Rβ1 resulted in increased dissemination of bacteria to the periphery and reduced ability to generate IFN-γ producing T cells ^15, 18^. We wanted to determine whether this observation was unique to the resistant B6 strain of mouse or if there was differential expression in mice which are substantially more susceptible to the low dose aerosol challenge^38^. To achieve this we infected resistant B6 and susceptible CBA/J (CBA) and C3H/HeJ (C3H) mice in the same aerosol chamber and determined bacterial burden (Fig 1A) and expression of the full length and ΔTM-IL-12Rβ1 (Fig 1B) at day 15 post infection. We found that while the bacterial burden in the lung was not markedly dissimilar between the strains of mice at this time point there was a much greater range of dissemination to the mediastinal lymph node (MLN) and that there was no detectable dissemination to the spleen or liver in the C3H and CBA mice (Fig 1A). When we measured the copy number of both full length and ΔTM-IL-12Rβ1 mRNA by quantitative RT-PCR we found that while all three strains expressed a small but measurable amount of full length mRNA, the C3H and CBA mice made a much greater amount of ΔTM-IL-12Rβ1 in their lungs than did the B6. While there was modest induction of the ΔTM-IL-12Rβ1, the baseline level of mRNA for ΔTM-IL-12Rβ1 was already high in both the CBA and C3H suggesting that expression may be regulated by a homeostatic factor unrelated to Mtb infection. Regardless of the mechanism that drives baseline induction of ΔTM-IL-12Rβ1, it is clear that a high level of expression is associated with reduced dissemination of bacteria from the lung to the periphery.

**Figure 1.**
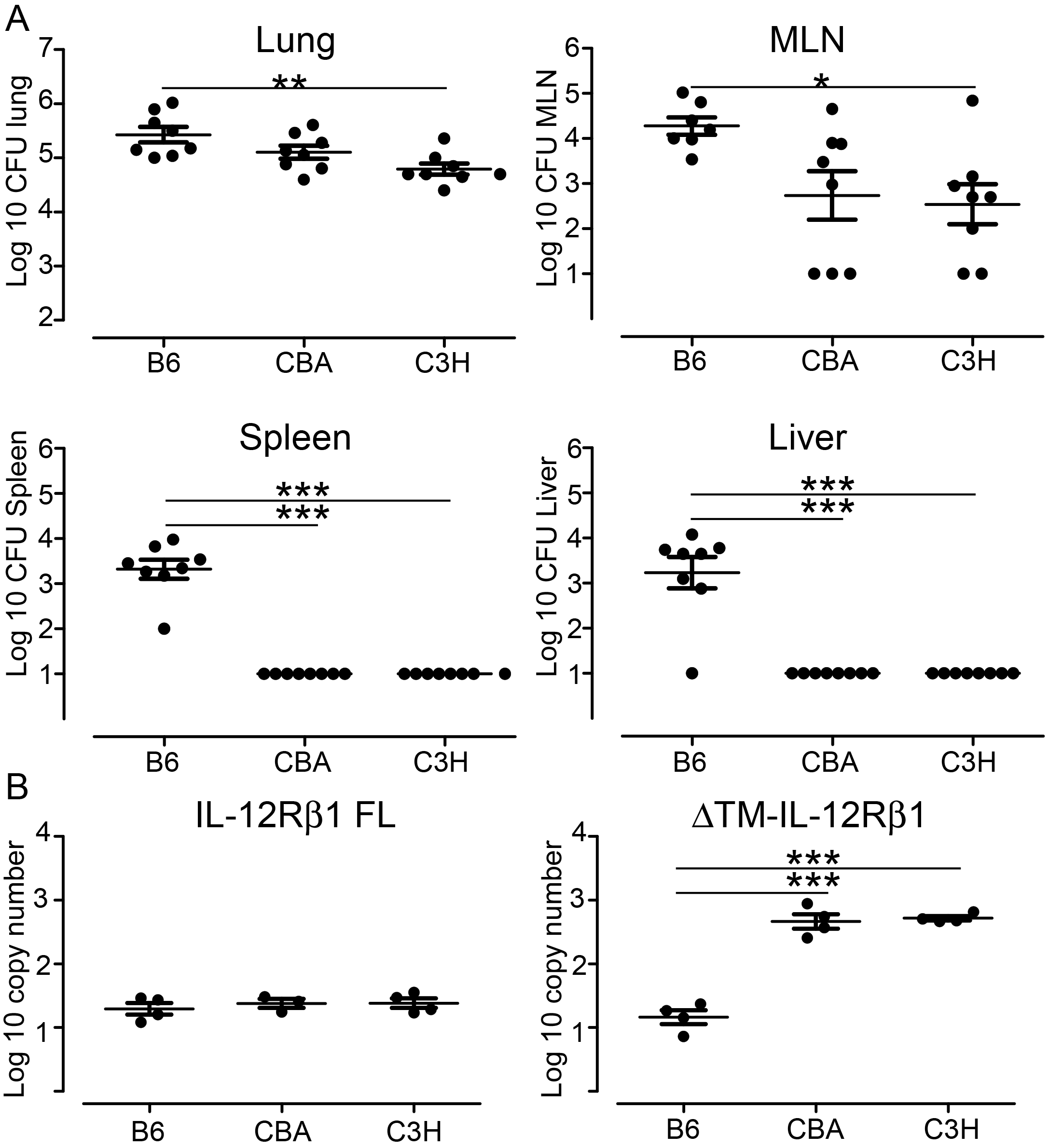
Mice susceptible to Mtb infection express high levels of ΔTM-IL-12Rβ1 and limit early dissemination form the lung to the periphery. Mice were infected using an aerosol chamber capable of reproducibly delivering approximately 75 colony forming units to each mouse. Mice were housed for 15 days after challenge and then the organs plated for determination of colony forming units (CFU) (Fig 1A) and the copy number of mRNA for the full length IL-12Rβ1 (FL-IL-12Rβ1) and the alternative splice variant ΔTM-IL-12Rβ1 determined by quantitative RT-PCR (Fig 1B). All data points are shown from 2 independent experiments (Fig 1A) or one data set representative of two total experiments (Fig 1B). Statistical significance was determined using one way ANOVA comparing all groups; n = 8 per group (Fig 1A) and n= 4 per group repeated twice (Fig 1B). *P<0.05, **P<0.01, ***P<0.001.

### Expression of ΔTM-IL-12Rβ1 can be induced in human dendritic cells and is impacted by TB in myeloid cells and lymphocytes in blood

Having identified an association between high level expression of ΔTM-IL-12Rβ1 and reduced dissemination in mice with increased susceptibility to TB we wanted to determine expression of ΔTM-IL-12Rβ1 in specific cells in humans with and without TB. In a first study we extended our observation that human monocyte-derived dendritic cells express ΔTM-IL-12Rβ1 in response to Mtb^8^ by determining the kinetics of the response to live Mtb exposure in monocyte-derived dendritic cells from several anonymous blood donors. By measuring the copy number of full length and ΔTM-IL-12Rβ1 mRNA using quantitative RT-PCR we were able to generate a ratio between the two signals and compare the response of individual donors. We found that donors made a strong ΔTM-IL-12Rβ1 response between 3 and 6 hours which waned by 9 hours (Fig 2A). These data suggest that Mtb drives an acute but not prolonged expression of ΔTM-IL-12Rβ1 in this cell type. We also probed the gene-expression array data in monocytes, neutrophils and lymphocytes that had been isolated from the blood of healthy control and those with active pulmonary TB^31^. We found that the signal from the probe specific for the ΔTM-IL-12Rβ1 was more highly expressed in monocytes and neutrophils from TB patients relative to controls (Fig 2B) and that while the CD4 and CD8 T cell signals independently failed to achieve the statistical threshold for significance, when the data sets were combined and analyzed as T lymphocytes, they were significantly different (average log2 intensity: PTB 8.145+/−0.10, n=14 versus HC 7.72+/−0.16, n=8; P=0.0294). These data indicate that exposure to Mtb and active TB induces expression of ΔTM-IL-12Rβ1 in human cells of both myeloid and lymphoid origin.

**Figure 2.**
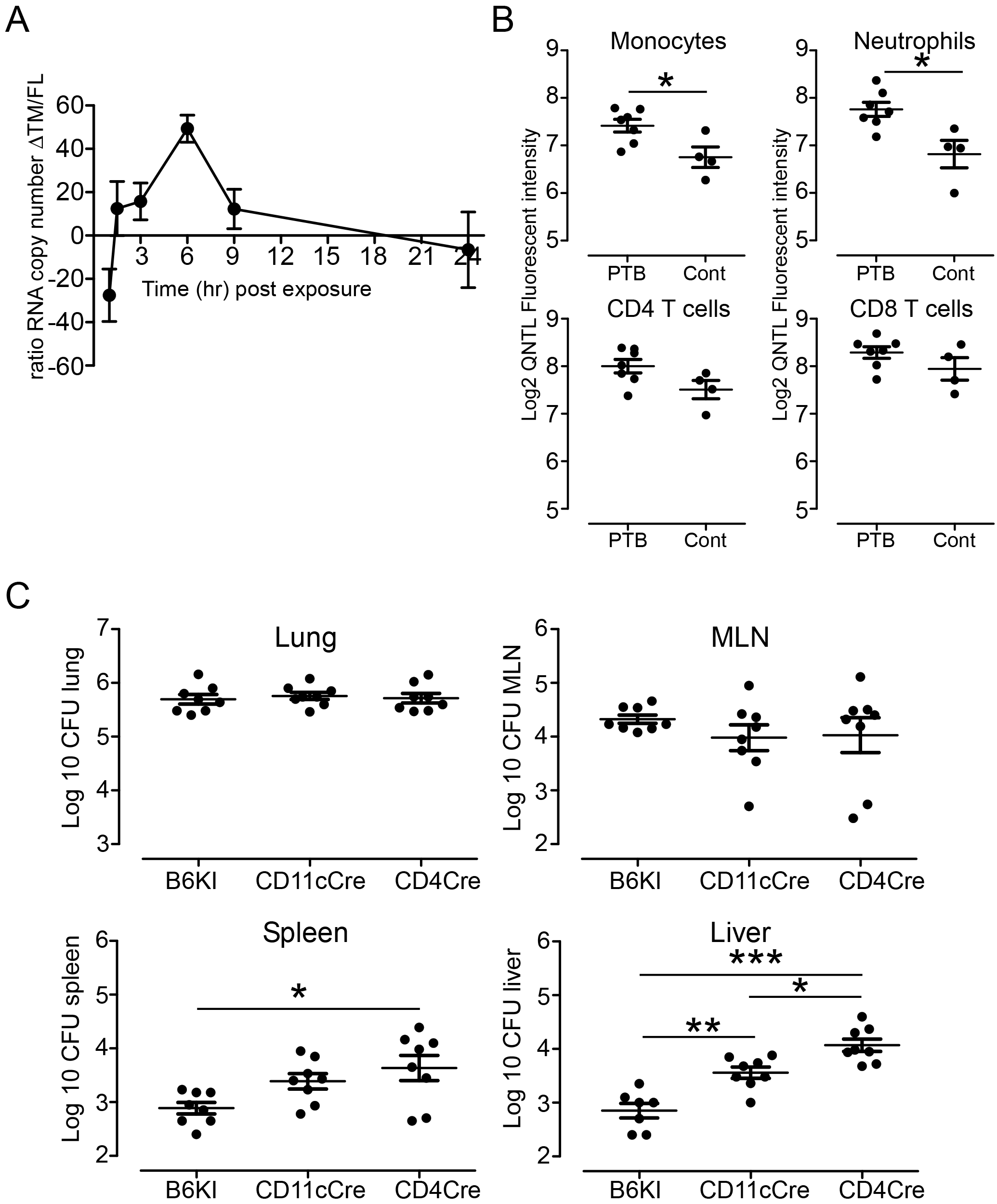
ΔTM-IL-12Rβ1 is expressed by lymphoid and myeloid human cells and its absence in CD11c or CD4 expressing cells compromises control of dissemination in mice. Human myeloid derived dendritic cells were generated from anonymous blood donors and exposed to live Mtb for 24 hours. RNA was extracted over the time course and analyzed for the copy number of mRNA for the full length IL-12Rβ1 (FL-IL-12Rβ1) and the alternative splice variant ΔTM-IL-12Rβ1 by quantitative RT-PCR. The ratio of ΔTM-IL-12Rβ1 to FL-IL-12Rβ1 was determined for each donor and the data points represent the mean of 4-7 values per time point (Fig 2A). Array data from GSE19443 was analyzed for expression of the ΔTM-IL-12Rβ1 in cells purified from the blood of healthy controls (Cont) and TB patients (PTB). Data points are all shown with an n=4-7, the difference between the means was determined by Students *t* test, * P<0.05 (Fig 2B). Mice either intact (B6KI) lacking the ability to generate the ΔTM-IL-12Rβ1 in either CD11c (CD11cCre) or CD4 (CD4Cre) expressing cells were infected with Mtb and the bacterial burden determined at day 15 as described for Fig 1. All data points are shown for 2 independent experiments, n=8, differences between the mean were determined by ANOVA with all columns being compared. *P<0.05, **P<0.01, ***P<0.001 (Fig 2C).

### Inability of either CD11c or CD4 expressing cells to produce ΔTM-IL-12Rβ1 results in increased dissemination of Mtb from the lung to the periphery following aerosol infection

We see expression of the ΔTM-IL-12Rβ1 in both myeloid and lymphoid cells in blood from TB patients (Fig 2B) and we know that a global inability to make ΔTM-IL-12Rβ1 reduces control of dissemination^15^ while high expression is associated with inhibition of dissemination of Mtb (Fig 1). We wanted therefore to determine whether myeloid or lymphoid cell expression of ΔTM-IL-12Rβ1 is responsible for the control of dissemination of bacteria from the lung to the periphery. To do this we crossed our transgenic mouse, which excises exons 13 and 14 in a FLP/CRE dependent manner resulting in expression of the full length IL-12Rβ1 without the structure to splice the ΔTM-IL-12Rβ1^15^ to mice expressing Cre under the CD11c or CD4 promoter. We infected the mice via aerosol and determined bacterial burden in the lungs and periphery at day 15 post infection (Fig 2C). We found that while the bacterial burden in the lung was not affected by cell specific loss of ΔTM-IL-12Rβ1 expression, there was a significant increase in the bacterial burden in the liver of mice lacking ΔTM-IL-12Rβ1 in both CD11c and CD4 expressing cells (Fig 2C). While the mean number of bacteria in the MLN was not significantly different, the range of bacterial numbers was much increased for the mice lacking ΔTM-IL-12Rβ1 in both CD11c-and CD4-expressing cells. Further, while significance was seen for increased dissemination to the spleen in mice lacking ΔTM-IL-12Rβ1 in CD4-expressing cells, there was a trend to increased dissemination in the mice where ΔTM-IL-12Rβ1 was deleted in CD11c^+^ cells. These data support the hypothesis that cells of both the lymphoid and myeloid lineage produce ΔTM-IL-12Rβ1 in response to infection and that while the lymphoid compartment seems to mediate the strongest effect, there is a measurable role for myeloid-derived ΔTM-IL-12Rβ1 in TB control. In preliminary data we find that absence of ΔTM-IL-12Rβ1 in CD11c expressing cells results in disruption of cytokine balance in the lung resulting in more IL-12p40 and more IL-10 (data not shown).

### Meta-analysis of transcriptomic data reveals an association between PTB and expression of the ΔTM-IL-12Rβ1 in human blood

We observe that Mtb induces ΔTM-IL-12Rβ1 expression in human cells (Fig 2A) and that purified cells from the blood of TB patients express ΔTM-IL-12Rβ1 (Fig 2B). We also observe an association between ΔTM-IL-12Rβ1 expression and dissemination from the lung to the periphery in the experimental mouse model (Fig 1 and Fig 2C). Together the data support analysis of ΔTM-IL-12Rβ1 expression in human disease. To achieve this we used defined criteria to select publically available array data sets and obtained 10 data sets from 5 countries, 3 continents and 4 platforms (Table 2) ^24, 31, 39, 40, 41^. In both Illumina and Affymetrix array platforms there are 3 probes representing detection of IL12RB1 transcription, one of which is specific for the full length transcript, one specific for the ΔTM-IL-12Rβ1 and one that can detect both. To determine what each of the probes contributes to the observed total IL12RB1 signal, we analyzed the raw data for each probe in 4 array data sets (Fig 3). This analysis revealed that for more than 90% of the samples, the signal from the probe specific for the full length IL-12Rβ1 transcript is negligible (i.e. the raw probe signal is insignificant) (Fig 3, FL-IL12Rβ1) while the signal for ΔTM-IL-12Rβ1 is detectable in almost all samples (Fig 3, ΔTM-IL-12Rβ1). The intensity of the signal for the probe specific for ΔTM-IL-12Rβ1 is highly correlated with the signal from the probe which detects both the full length and ΔTM-IL-12Rβ1 transcripts, while the signal for the probe specific for the full length transcript correlates with both signals to a much lower degree (Table 3). These analyses suggest that blood cells express the ΔTM-IL-12Rβ1 transcript in significant amounts and that this transcript is the dominant one.

**Table 3.** The signal from probes specific for the ΔTM-IL-12Rβ1 isoform in microarray data sets correlates with the majority of signal seen for all of the IL-12Rβ1 targeted probes.

| Array data set | Correlation coefficient between the signal for probes specific for: |  |  |
| --- | --- | --- | --- |
| | ΔTM-IL-12Rβ1 and all probes | ΔTM-IL-12Rβ1 and IL-12Rβ1 full-length | IL-12Rβ1 full-length and all probes |
| GSE19444 | 0.66<sup>1</sup> | 0.43 | 0.32 |
| GSE19442 | 0.44 | -0.16 | -0.09 |
| GSE56153 | 0.89 | 0.67 | 0.68 |
| GSE54992 | 0.51 | 0.16 | 0.05 |

<sup>1</sup> Normalised fluorescent intensities of probes specific for different isoforms of IL-12Rβ1 were used to prepare a matrix and the Pearson correlation coefficient was calculated using online tool Metaboanalyst 3.0.

**Figure 3.**
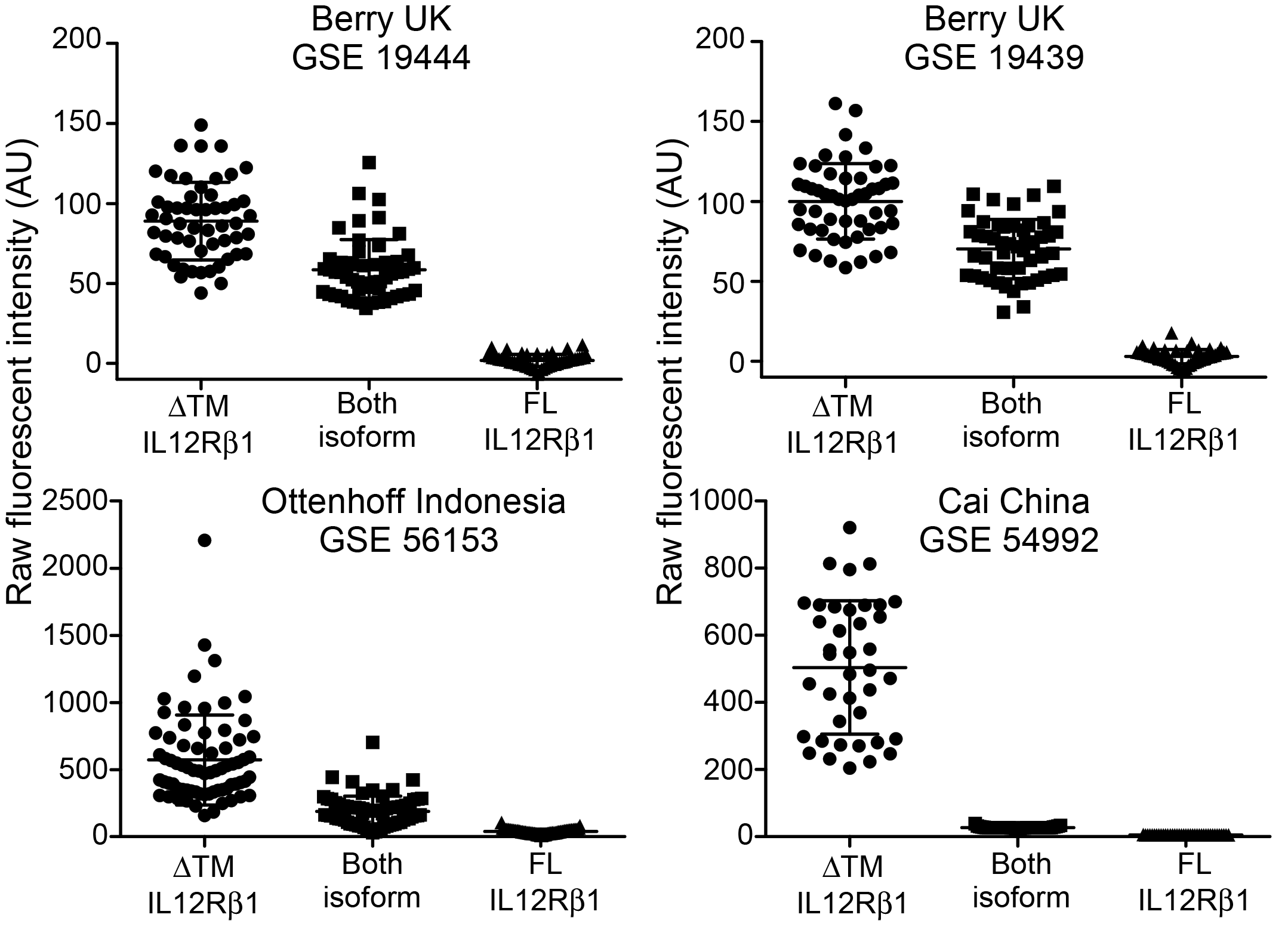
The majority of the IL-12Rβ1 signal seen in microarray data is derived from the ΔTM-IL-12Rβ1 specific probe. Raw data was manually curated and the signal for each probe analyzed separately for each of 4 microarray data sets. The raw fluorescent intensity for each sample and different IL-12Rβ1 isoform specific probe is shown. The probe intensity for the FL-IL-12Rβ1 was not considered to be significant for the majority of samples and over all of the data sets whereas the signal from the probe specific for the ΔTM-IL-12Rβ1 was significantly expressed in all samples.

### Higher expression of the ΔTM-IL-12Rβ1 transcript correlates with pulmonary TB

We see a specific signal for ΔTM-IL-12Rβ1 in transcriptional data sets from human blood (Fig 3) and we know that this molecule is associated with altered disease outcomes in the experimental mouse model (Fig 1 and 2). These observations support an in depth analysis of the transcriptional data sets from individuals with pulmonary TB and their controls. To achieve this we processed, transformed and normalized the raw data from the transcriptional arrays identified in Table 2 and compared the signal to the sample groups defined by the investigators originating the data sets. The signal for the probe specific for ΔTM-IL-12Rβ1 significantly differentiated PTB and control groups in all arrays tested in the UK^31^, China^24^, Indonesia^40^, Malawi^39^ and South Africa^39^ (Figure 4A). One data set in South Africa showed a trend but did not quite make significance (GSE19442)^31^ and the data in Indonesia^40^ had a very small difference as might be expected as the only difference between the groups was the absence of active symptoms for TB (i.e. non TB rather than healthy control). Despite the small difference the data was not variable and thus reached significance statistically. These data demonstrate that ΔTM-IL-12Rβ1 expression in the blood is associated with active pulmonary TB.

**Figure 4.**
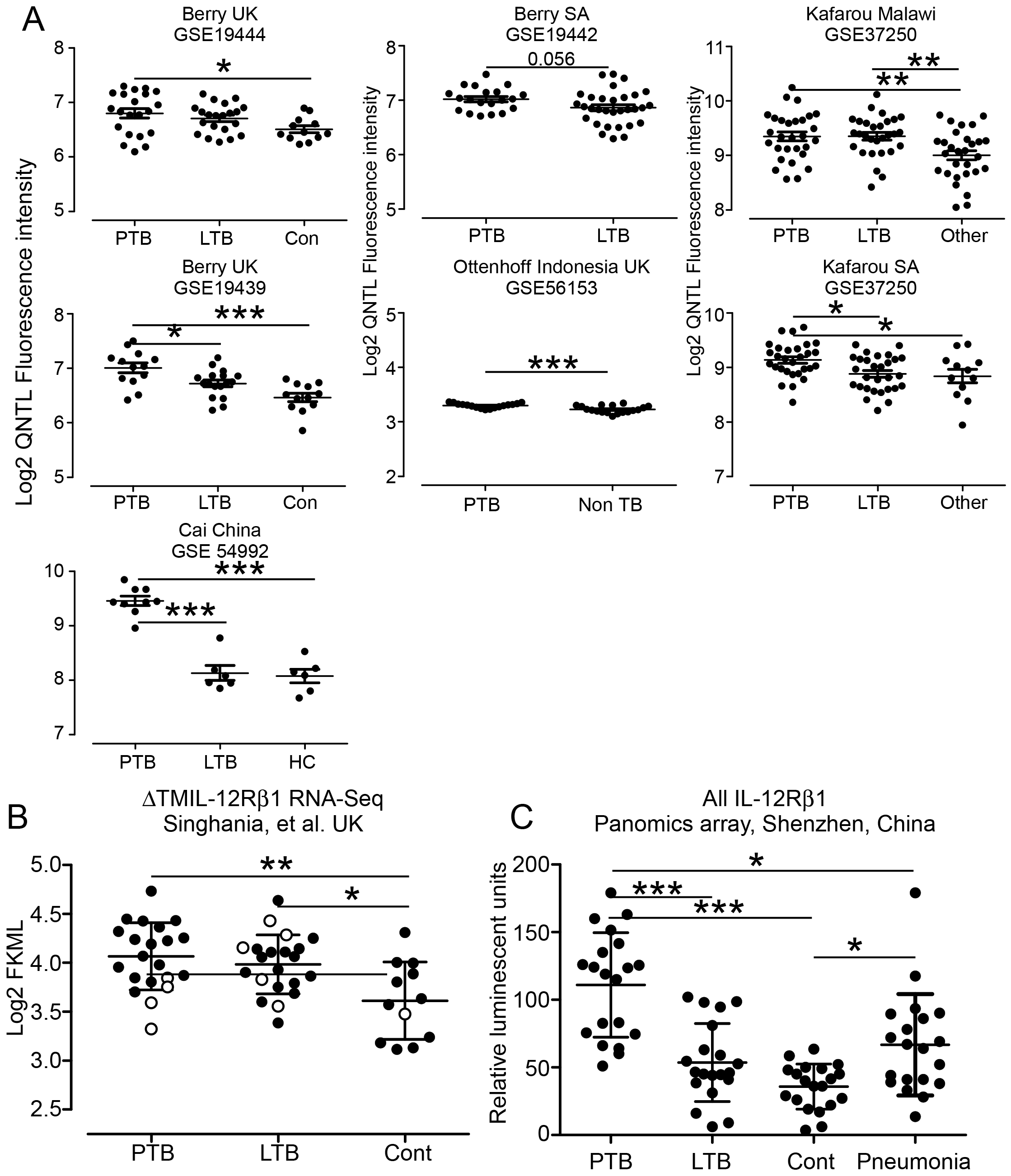
Expression of ΔTM-IL-12Rβ1 in blood associates with active pulmonary TB and is variably associated with latent TB. Microarray data from publically available databases was analyzed for the signal specific for ΔTM-IL-12Rβ1 (Fig 4A). All data points are shown and the differences between the means of the ΔTM-IL-12Rβ1 signal for each group determined by ANOVA with Tukey’s multiple comparison, n=6-31, *P<0.05, **P<0.01, ***P<0.001. RNA-Seq data sets from a UK cohort was analyzed for expression of the ΔTM-IL-12Rβ1 (Fig 4B) (all data points are shown with known group outliers shown as open symbols) and was discriminatory for controls versus both LTB and PTB, using ANOVA with Tukey’s multiple comparison, n=12-21, *P<0.05, **P<0.01. RNA was taken from a new cohort in China and analyzed for expression of all IL-12Rβ1 transcripts and was shown to discriminate between controls and LTB and PTB, between LTB and PTB and between PTB and pneumonia, using ANOVA with Tukey’s multiple comparison, all data points are shown, n=20, *P<0.05, ***P<0.001.

### Expression of the splice variant in the blood reflects the heterogeneity of latent TB infection

While the ΔTM-IL-12Rβ1 transcript clearly associates with pulmonary TB (PTB), it is variably able to discriminate between latent TB infection (LTB) and PTB (Fig 4A) with 3 of 7 studies showing discrimination but no association with location. One study showed very clear discrimination (China GSE 54992)^24^ and this may reflect the fact that this resulted from isolated PBMC rather than whole blood or may just reflect the difference in the platform. These data suggest that the ΔTM-IL-12Rβ1 may have the capacity to discriminate between LTB and PTB but work needs to be done to improve the signal. Within the three studies which showed the ability to discriminate between LTB and PTB we investigated whether the ΔTM-IL-12Rβ1 signal was the strongest IL-12-related signal capable of discriminating between infection outcomes. To do this we generated a data matrix using array signals from IL12A, IL12B, IL12RB1 and IL-12RB2 as well as ratios of each set of signals as variables to see how well they differentiated between groups. The top three performing variables based on area under the curve (AUC) or receiver operating characteristic (ROC) curve capable of discriminating between control and PTB, LTB and PTB and LTB and control are shown in Table 4. The signal for ΔTM-IL-12Rβ1 either alone or as a ratio with other members of the IL-12 family appears in the top three for all three arrays and for all three discriminatory activities. These analyses demonstrate that the ΔTM-IL-12Rβ1 signal has the potential to be a diagnostic tool and should be studied further.

**Table 4.**
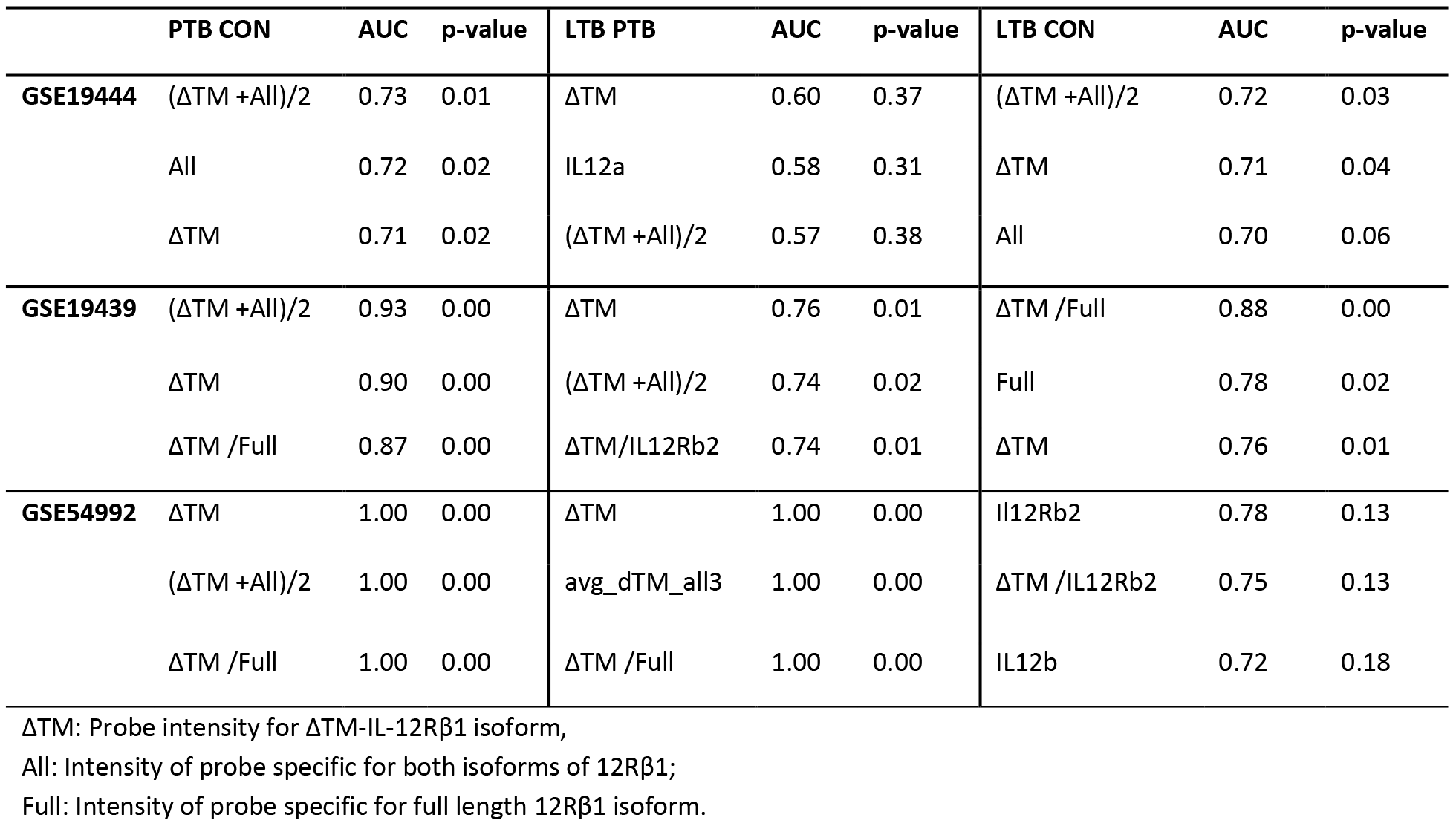
Discriminatory potential of IL-12-related variables for identifying disease groups. Variables which include the signal from the ΔTM-IL-12Rβ1 occur as one of the top 3 variables IL-12-related discriminator in each comparison. The top 3 variables based on area under curve (AUC) for each comparison is shown.

To extend our analysis of array data we analyzed RNA-Seq data generated from the RNA used in the GSE19444 UK^31,30^ for transcription of ΔTM-IL-12Rβ1 and found that as for array data, the ΔTM-IL-12Rβ1 signal discriminates between PTB and controls, the ΔTM-IL-12Rβ1 signal also discriminates between LTB and controls (Fig 4B). Importantly, we identify those samples identified as outliers in previous analyses,^31,30^ by open circles (Fig 4B) which show low expression of ΔTM-IL-12Rβ1 in the PTB outliers and high expression in 3 of 5 in the LTB outliers suggesting that the ΔTM-IL-12Rβ1 signal may be associated with the discriminatory gene-signature identified previously^31,30^.

Finally, we wanted to test the predictive and discriminatory power of the ΔTM-IL-12Rβ1 signal in a test patient cohort in China. To do this we probed the RNA from whole human blood using a targeted array capable of measuring the IL-12Rβ1 signal. We found that while the LTB cohort had a more variable signal than the controls, there was no significant difference between the means for these two populations and the IL-12Rβ1-specific signal discriminated effectively between control and PTB, LTB and PTB, PTB and pneumonia (Fig 4C). These data further support the utility of the ΔTM-IL-12Rβ1 signal to discriminate between PTB and LTB and PTB and other diseases.

### The ΔTM-IL-12Rβ1 signal decreases upon treatment in areas of low infection pressure

Because we see a difference in the ΔTM-IL-12Rβ1 signal between controls and PTB we reasoned that the ΔTM-IL-12Rβ1 may provide a signal indicating that treatment is effective. To determine if this was the case, we analyzed the transcriptome sets containing data from treated individuals from areas of low infection pressure (UK, overall incidence rate 10/100,000 – but with pockets of moderate infection pressure), moderate infection pressure (China, incidence rate 64/100,000) and high infection pressure (South Africa, incidence rate 781/100,000). We found that in the UK and China an early (8-12 weeks) drop in the ΔTM-IL-12Rβ1 signal was seen during treatment, which was sustained long term (Fig 5, top graphs). In contrast, in two studies in South Africa no sustainable or reproducible drop in signal was seen over the population during treatment (Fig 5, bottom graphs). These data suggest that the signal for ΔTM-IL-12Rβ1 may be useful to detect treatment success in situations where the risk of reinfection is low, but it may reflect repeated infection where reinfection risk is high. Determining when and how the ΔTM-IL-12Rβ1 is expressed in the lungs of those recently exposed to TB is therefore an important undertaking.

**Figure 5.**
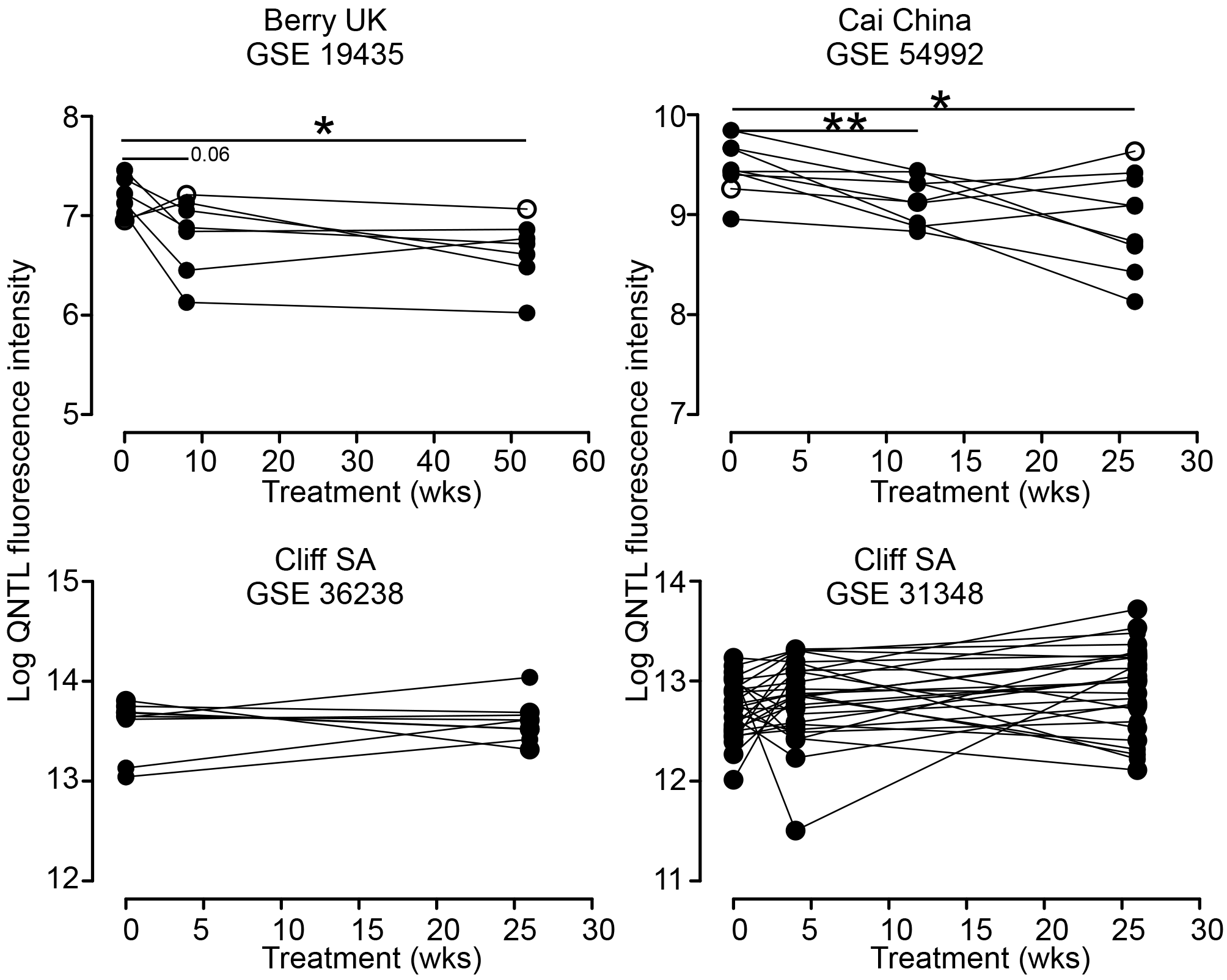
Expression of ΔTM-IL-12Rβ1 in blood associates drug treatment in areas of low to moderate infection pressure but fails in areas where infection pressure is high. Publically available array data was analyzed for the expression of ΔTM-IL-12Rβ1 in the blood of TB-infected individuals undergoing drug treatment. All data points are shown and the ability of the signal to differentiate between pretreatment and a short and longer period post treatment determined. Paired *t* test was used to compare values at 0 and 6 or 8 week and 0 and 26 or 52 weeks for each patient. In the UK and China (top graphs) significant differences in the means were observed after treatment whereas no discernable effect could be seen in the South African cohorts undergoing treatment, n=7-27 P<0.05, **P<0.01.

## Discussion

We show here that the ΔTM-IL-12Rβ1 is a significant indicator of active TB in human populations across the globe, that it can discriminate between latent and active TB under some conditions and that it drops during drug treatment in areas where the risk of reinfection is low. These observations are valuable and suggest that this gene transcription signal can contribute to the development of sensitive and specific modular transcriptional signatures ^30^ capable of defining the risk of TB development in those infected with the bacterium. The goal for these signatures is to eventually turn the marked heterogeneity of latent infection from a black box into a fully defined continuum with distinct transcriptional markers capable of allotting individuals into low, mid and high risk of progression. In addition to this goal however the data we report here also highlight the potential for the ΔTM-IL-12Rβ1 signal to inform us about the mechanisms underlying the development and control of TB in the human lung.

In considering the early events in the TB-exposed lung, the invading bacteria is unlikely to meet any activated macrophages or T cells when it first arrives and thus for the acquired immune response to be initiated, the bacteria must migrate to the draining lymph node^12^. This migration is slow to occur and this delay in activation and arrival of T cells allows the bacteria to grow unrestrained for up to 2 weeks in the B6 mouse^9, 42^. Cytokines and chemokines capable of initiating early responses are therefore key to the rapidity of the response and to eventual outcome^43^. We know that IL-12p40 is expressed very early following Mtb infection and that its production precedes production of IL-23 or IL-12p35 suggesting that it functions alone ^7, 8^. We also know that type 1 IFN, IL-1, IL-10 and IL-12 cross regulate each other in response to Mtb infection both in vitro and in vivo ^7 44, 45, 46^ and that the relative level of these cytokines at the initiation of infection has the potential to impact the initiation of immunity. Our data reported here suggest that the very high expression of the ΔTM-IL-12Rβ1 in the C3H and CBA mice may sequester IL-12p40, which we know aids in overcoming the IL-10-induced resistance to migration in Mtb-exposed dendritic cells ^7^. This inhibition may then severely limits the migration of bacteria from the lung to the periphery – as is seen in the CBA/J and C3H mice. In turn this delay then results in slow induction of T cells thus leading to the observed higher susceptibility of these mice strains to Mtb ^38^. In contrast, total lack of the splice variant results in slightly higher dissemination of bacteria from the lung ^15^ and this effect can be seen if either myeloid or lymphoid cells lack the ΔTM-IL-12Rβ1. The increased effect of the absence of the ΔTM-IL-12Rβ1 in CD4 cells may reflect the role of the ΔTM-IL-12Rβ1 in augmenting IL-12p70-induced IFN-I production in T cells^15, 18^, while the impact of the loss of ΔTM-IL-12Rβ1 in the CD11c expressing cells may reflect the disruption of protective cytokine balance at the beginning of infection. The very early but short lived ΔTM-IL-12Rβ1 response of the human dendritic cells to live Mtb infection suggests that the signal is highly regulated and that sustained expression in myeloid cells may reflect ongoing exposure to live bacteria. In this regard, our data showing increased expression of ΔTM-IL-12Rβ1 signal in the blood of active TB patients suggests that myeloid cells expressing this signature may have been exposed to live bacteria in the recent past. Similarly, the drop in ΔTM-IL-12Rβ1 signal seen during drug treatment may reflect removal of live bacteria from the body and therefore the loss of stimulation for the signal. If one considers the infection pressure in the areas where the ΔTM-IL-12Rβ1 signal drops following drug treatment, it is possible that there is little re-exposure to drive the signal up again but in areas where re-infection is likely there will be repeated exposures in the lung and the ΔTM-IL-12Rβ1 will be maintained despite drug treatment.

IL-12Rβ1 exhibits a marked degree of variability at the genomic5, transcriptional^47^ and splice variant level^17^ suggesting that it is a highly flexible receptor with the capacity to both stimulate and inhibit cellular responses. Our observation that ΔTM-IL-12Rβ1 is secreted^15^ suggests that it acts to sequester cytokine and regulate activation of cells however it appears to drive activation of T cells ^8, 18^ and is a positive regulator of immunity. Its effects in mouse lungs suggests that, just as for type 1 IFN^48^ a little bit is good and required for optimal control of bacterial dissemination but too much completely limits dissemination and hampers induction and expression of protective immunity.

Our data demonstrate that determining the underlying mechanisms whereby the alternative splice variant of IL-12Rβ1 impacts TB immunity in both mouse and man should be pursued. The data also support further investigation into the best use of the transcriptional ΔTM-IL-12Rβ1 signal in defining the heterogeneity of latent TB infection.

